# Xylazine effects on opioid-induced brain hypoxia

**DOI:** 10.1101/2023.03.31.535103

**Authors:** Shinbe Choi, Matthew R. Irwin, Eugene A. Kiyatkin

## Abstract

Xylazine has emerged in recent years as an adulterant in an increasing number of opioid-positive overdose deaths in the United States. Although its exact role in opioid-induced overdose deaths is largely unknown, xylazine is known to depress vital functions and cause hypotension, bradycardia, hypothermia, and respiratory depression. In this study, we examined the brain-specific hypothermic and hypoxic effects of xylazine and its mixtures with fentanyl and heroin in freely moving rats. In the temperature experiment, we found that intravenous xylazine at low, human-relevant doses (0.33, 1.0, 3.0 mg/kg) dose-dependently decreases locomotor activity and induces modest but prolonged brain and body hypothermia. In the electrochemical experiment, we found that xylazine at the same doses dose-dependently decreases nucleus accumbens oxygenation. In contrast to relatively weak and prolonged decreases induced by xylazine, intravenous fentanyl (20 μg/kg) and heroin (600 μg/kg) induce stronger biphasic brain oxygen responses, with the initial rapid and strong decrease, resulting from respiratory depression, followed by a slower, more prolonged increase reflecting a post-hypoxic compensatory phase, with fentanyl acting much quicker than heroin. The xylazine-fentanyl mixture eliminated the hyperoxic phase of oxygen response and prolonged brain hypoxia, suggesting xylazine-induced attenuation of the brain’s compensatory mechanisms to counteract brain hypoxia. The xylazine-heroin mixture strongly potentiated the initial oxygen decrease, and the pattern lacked the hyperoxic portion of the biphasic oxygen response, suggesting more robust and prolonged brain hypoxia. These findings suggest that xylazine exacerbates the life-threatening effects of opioids, proposing worsened brain hypoxia as the mechanism contributing to xylazine-positive opioid-overdose deaths.

## Introduction

A new player in the US opioid epidemic is xylazine, which is a non-controlled substance traditionally used as a veterinary tranquilizer and component of general anesthesia in animals [1]. Although xylazine is not approved for human use, a pattern of recreational use in the US has emerged in the past decade [2]. Reports from the Drug Enforcement Administration show that xylazine-positive fatal overdoses have experienced a significant jump from 2020 to 2021, especially in the South region where the noted increase was 1127% [3]. More recently, in 2022, xylazine was found as an adulterant in a significant percentage of opioid-related overdose deaths, appearing in 25.8% of total overdose deaths in Philadelphia and 19.3% of total overdose deaths in Maryland [4]. Fentanyl was present in 98% and heroin in 23% of xylazine-involved deaths, revealing a dangerous connection between xylazine and these opioid drugs.

Xylazine’s exact role in opioid-induced overdose deaths is largely unknown. Similar to clonidine, xylazine is an agonist at the alpha-2 adrenergic receptors that induces sedation and muscle relaxation [5]. At higher doses, xylazine has been shown to significantly depress vital functions, causing strong hypotension, bradycardia, hypothermia, and respiratory depression [6]. When taken with opioids that share many common effects with xylazine, the risk of overdose and death may increase. However, the mechanisms underlying the effects of xylazine and its interaction with opioid drugs are still relatively unknown.

In this study, we focused on the effects of xylazine as an adulterant drug taken in combination with heroin and fentanyl. These two highly potent opioid drugs induce strong respiratory depression and robust brain hypoxia at low doses [7–10]. To learn more about the basic physiological and behavioral effects of xylazine, we used multi-site thermorecording coupled with monitoring of locomotor activity. As temperature is an important homeostatic parameter, simultaneous recordings from the brain site, temporal muscle and subcutaneous space provides a valuable tool for clarifying the mechanisms underlying changes in brain temperature and assessing the metabolic and vascular effects of xylazine [11].

Since xylazine is a CNS depressant that can inhibit respiration at high doses, we used oxygen sensors coupled with high-speed amperometry to examine the effects of xylazine on brain oxygenation. Although the hypoxic effects of drugs can be assessed by plethysmography [7,12,13] or pulse oximetry [14–16], it is unclear how changes in breathing activity or hemoglobin saturation in blood translate into changes in oxygen levels in the brain’s extracellular space—a functionally important parameter that affects the health and survival of neural cells. In contrast to these peripheral measures, oxygen sensors allow direct assessment of real-time fluctuations in brain oxygenation induced by natural stimuli, occurring during motivated behavior, and after exposure of different drugs in freely moving rats under physiologically relevant conditions [11, 17–21].

After we established the basic physiological effects of xylazine, we examined how this drug at a moderate, human-relevant dose affects changes in brain oxygenation induced by fentanyl and heroin. In these tests we compared changes in brain oxygen levels induced by fentanyl and heroin with their combined administration with xylazine. Like in our previous studies, the nucleus accumbens (NAc), a deep brain structure involved in sensorimotor integration and functioning of the motivation-reinforcement circuits [22–24], was the recording site in both thermorecording and electrochemical experiments.

## Materials and Methods

### Subjects

15 adult male Long-Evans rats (Charles River Laboratories) weighing 450±50 g at the time of surgery were used in this study. Rats were individually housed in a climate-controlled animal colony maintained on a 12-12 hr light-dark cycle with food and water available *ad libitum*. All procedures were approved by the NIDA-IRP Animal Care and Use Committee and complied with the Guide for the Care and Use of Laboratory Animals (NIH, Publication 865-23). Maximal care was taken to minimize the number of experimental animals and any possible discomfort or suffering at all stages of the study.

### Overview of the study

This study combines data obtained with two different technologies used in freely moving rats. By using multi-site thermorecording combined with monitoring of conventional locomotion, we assessed the basic behavioral, metabolic, and peripheral vascular effects of xylazine at doses within a range of possible human consumption. To assess drug-induced fluctuations in brain oxygenation, we used oxygen sensors coupled with high-speed amperometry. First, we examined the effects of xylazine at different doses on brain oxygenation. Second, we examined the changes in brain oxygenation induced by a fentanyl-xylazine mixture and compared them with the effects of fentanyl alone. To assess whether the effects of xylazine-fentanyl mixture are generalized to other opioid drugs, we examined the pattern of oxygen fluctuations induced by co-administration of xylazine with heroin and compared them with the effects of heroin alone.

### Surgical preparations

In both thermorecording and electrochemical experiments, we used similar surgical preparations described in detail elsewhere [25, 26]. Under general anesthesia (ketamine 80 mg/kg + xylazine 8 mg/kg with subsequent dosing), each rat was implanted with a jugular catheter. For thermorecording experiments, the rat was implanted with three copper-constantan thermocouple sensers in the NAc shell, temporal muscle, and subcutaneously along the nasal ridge with the tip ~15 mm anterior to bregma. Target coordinates of the recordings in the right NAc shell were: AP +1.2 mm, ML ±0.8 mm, and DV +7.2-7.6 mm from the skull surface, according to coordinates of the rat brain atlas [27]. For electrochemical experiments, each rat was implanted in the same NAc location with a Pt-Ir oxygen sensor (Model 7002-02; Pinnacle Technology, Inc., Lawrence, KS, USA). The probes were secured with dental cement to the three stainless steel screws threaded into the skull. In both experiments, the jugular catheter ran subcutaneously to the head mount and was secured to the same head assembly. Rats were allowed a minimum of 5 days of post-operative recovery and at least 3 daily habituation sessions (~6 h each) to the recording environment; jugular catheters were flushed daily with 0.2 ml heparinized saline to maintain patency.

### Electrochemical detection of oxygen

Pinnacle oxygen sensors consist of an epoxy-sheathed disc electrode that is grounded to a fine surface using a diamond-lapping disc. These sensors are prepared from a Pt-Ir wire 180 μm in diameter, with a sensing area of 0.025 mm^2^ at the tip. The active electrode is incorporated with an integrated Ag/AgCl reference electrode. Dissolved oxygen is reduced on the active surface of these sensors, which is held at a stable potential of −0.6 V versus the reference electrode, producing an amperometric current. The current from the sensor is relayed to a computer via a potentiostat (Model 3104, Pinnacle Technology) and recorded at 1-s intervals, using PAL software utility (Version 1.5.0, Pinnacle Technology).

Oxygen sensors were calibrated at 37°C by the manufacturer (Pinnacle Technology) according to a standard protocol described elsewhere [20]. The sensors produced incremental current changes with increases in oxygen concentrations within the wide range of previously reported brain oxygen concentrations (0-40 μM). Substrate sensitivity of each sensor varied from 0.57-1.19 nA/1μM. Oxygen sensors were also tested by the manufacturer for their selectivity toward other electroactive substances, including dopamine (0.4 μM) and ascorbate (250 μM), none of which had significant effects on reduction currents.

### Experimental procedures

A similar protocol was utilized in both thermorecording and electrochemical experiments. At the beginning of each experimental session, rats were minimally anesthetized (<2 min) with isoflurane and sensors (either thermocouple or oxygen) were connected via an electrically shielded flexible cable and a multi-channel electrical swivel to the recording instruments. The injection port of the jugular catheter on the head mount was connected to a plastic catheter extension that allowed stress and cue-free drug delivery from outside the cage. When the rats received two different drugs within one recording session, two catheter extensions mounted on the recording cable were used to minimize any contamination of one drug by another drug. Testing began a minimum of 90 min after connecting the sensors to the recording instruments, when baseline values of temperature and electrochemical currents stabilized. For the next 4-6 hours, rats received one of three drug treatments. Upon completion of drug treatments, rats were removed from the cages and briefly anesthetized by isoflurane to disconnect them from the recording instruments. Then, catheters were flushed with heparinized saline before rats were returned to the animal colony. Temperature recordings were combined with monitoring of locomotor activity using 4 infrared motion detectors (Med Associates) as previously described [25].

The recordings were conducted for several sessions (n=3-6), and the number of sessions in each experiment was determined by the quality of the recording and patency of the iv catheter over time. In the thermorecording experiment, we examined changes in temperatures and locomotion induced by iv xylazine (Xylazine hydrochloride, MP) at three doses (0.33, 1 and 3 mg/kg). These doses are lower than the generally accepted range of toxic effects of oral xylazine that vary in humans, between 40 and 2400 mg (or 0.6 and 34.3 mg /70 kg). The largest dose used (3 mg/kg) was well below the LD50 for iv xylazine in rats, which is between 22-43 mg/kg [28]. The drug was administered in an ascending order, with 60- and 90-min inter-injection intervals for the 0.33 and 1 mg/kg doses. Xylazine was delivered in 0.15-0.8 ml volumes of saline at a slow injection rate (0.2 ml/10 s).

Three types of tests were conducted in the electrochemical experiments. First, we mimicked the protocol of the thermorecording experiment and examined the effects of xylazine at the same three doses on NAc oxygenation. During the second treatment protocol, the rat received two iv injections of fentanyl (Fentanyl citrate injection 50 μg/mL; Fentanyl Citrate Injections; Hospira Inc.) at a 20 μg/kg dose both alone and as a mixture with xylazine at an effective dose determined in the first treatment protocol. As shown previously [9], 20 μg/kg is a modest dose that induces a biphasic brain oxygen response, with a rapid and strong decrease (to ~50% of baseline levels in drug-naïve rats) followed by more prolonged and weaker oxygen increase. In the third protocol, rats received two injections of heroin (Diamorphine Hydrochloride; obtained from NIDA-IRP Pharmacy) at a 600 μg/kg dose, both alone and as a mixture with xylazine at the same dose as in the second experiment. A 600 ug/kg dose is much larger than the optimal dose for heroin self-administration (75-100 μg/kg; [29]), but it induces a similar degree of decrease (~50% decrease) in NAc oxygen [30]. This dose difference is within the range of generally accepted differences in potencies of fentanyl and heroin to maintain iv self-administration behavior (ED50 2.5 ug/kg vs. 50 μg/kg or 1:20; [31]. In both the second and third treatment protocols, the second injection (xylazine-fentanyl or xylazine-heroin) was done 120 min after the first injection (fentanyl or heroin alone). The first and second drug protocols were conducted in the same rats and third protocol was conducted in a separate group of rats.

### Histological verification of electrode placements

After completion of the experiments, rats were sacrificed, and their brains were extracted and placed in 10% formalin. Brains were sliced on a cryostat and analyzed for verification of the location of cerebral implants as well as possible tissue damage around the recording site.

### Data analysis

Temperature data were sampled at 2-s time intervals and analyzed with 1-min time beans. They were presented as both absolute and relative changes with respect to the moment of drug administration. We also calculated NAc-muscle and skin-muscle temperature gradients that were used to determine the effects of xylazine on brain metabolic activity and tone of skin vasculature. Locomotor data were assessed with 1-min time resolution. Electrochemical data were sampled at 1 Hz using PAL software utility (Pinnacle Technology) and analyzed with 1-min and 10-s time resolutions. Electrochemical data were first analyzed as raw currents. Because each individual sensor differed slightly in background current and substrate sensitivity *in vitro*, currents were transformed into concentrations and represented as relative changes, with the pre-stimulus baseline set at 100%. One-way repeated measures ANOVAs (followed by Fisher LSD post-hoc tests) were used to evaluate statistical significance of drug-induced changes in temperature and brain oxygen changes. Two-way repeated-measure ANOVAs were used to analyze between-group differences in the effects of xylazine and its mixture with fentanyl and heroin.

## Results

### 1. Behavioral and temperature effects of xylazine

Xylazine, within the range of chosen doses, had sedative effects, decreasing temperature in all recording locations (Figure 1). Consistent with our previous studies [31], basal temperature was highest in the NAc, lower in temporal muscle, and lowest in the subcutaneous space (A). When analyzed as relative changes (B), temperature decreases were significant (see F values in Supplementary materials) and clearly dose-dependent. Despite parallel changes, the decrease was strongest in temporal muscle, weaker in the NAc, and lowest in the subcutaneous space. Due to these differences, NAc-muscle and skin-muscle temperature differentials significantly increased, with weaker changes for the former and much stronger changes for the latter (C). Figure 1D shows that xylazine tended to decrease locomotor activity; this effect was less evident at the lowest dose and more evident with higher drug doses.

**Figure 1.**
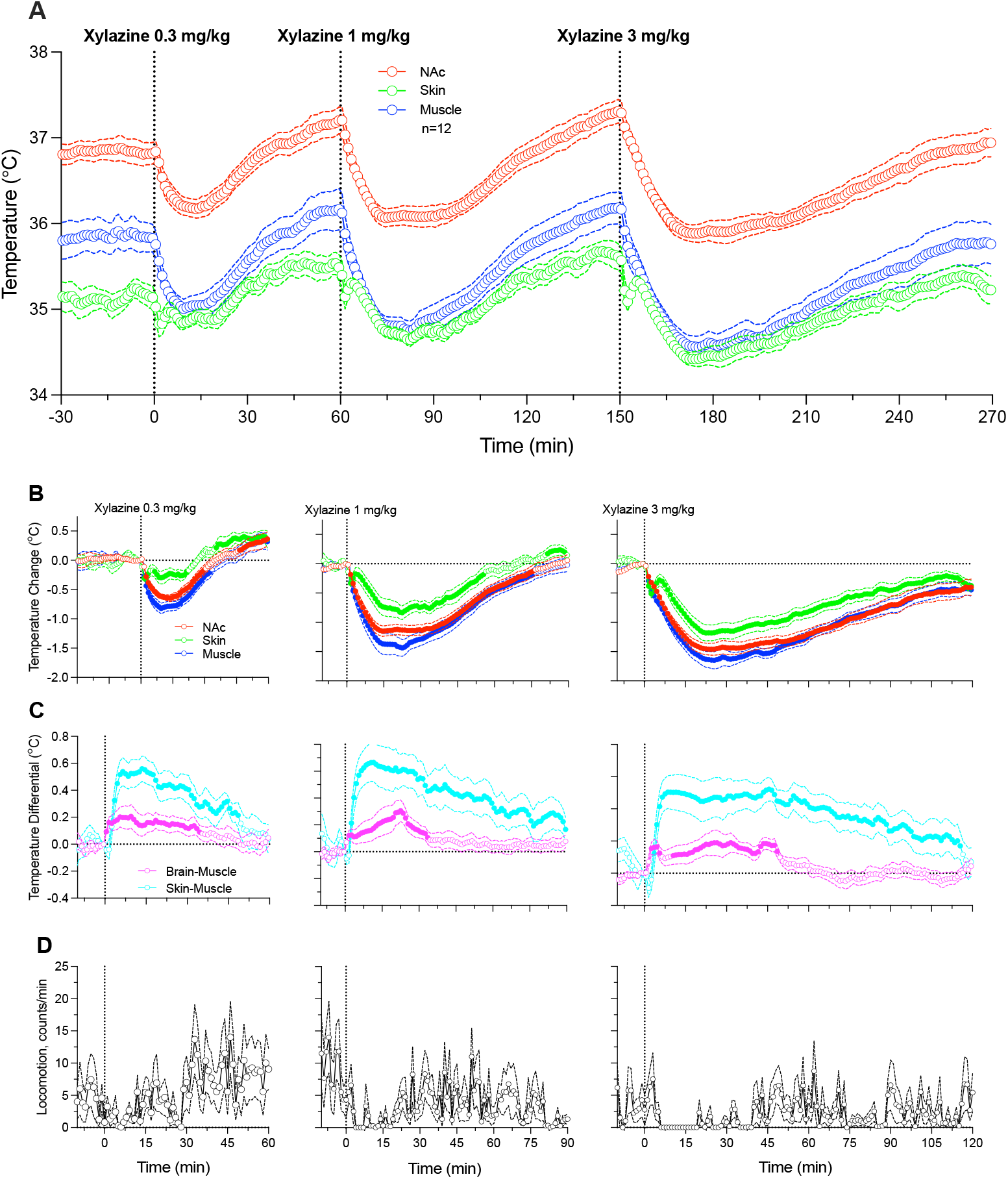
Changes in temperature induced by iv xylazine at different doses (0.33, 1.0 and 3.0 mg/kg in awake, freely moving rats. A = Absolute temperature changes. B = relative temperature changes; C = Brain-muscle and skin-muscle differentials. D = Locomotor activity. Filled symbols show values significantly different from pre-injection baseline.

### 2. Effects of xylazine on NAc oxygenation

The effects of xylazine on NAc oxygenation were examined in 5 rats during 11 daily sessions. As shown in Figure 2A, xylazine at each dose rapidly decreased NAc oxygen levels, then followed with a slower ascent to baseline. Xylazine-induced oxygen responses were clearly dose-dependent. At the lowest dose (0.33 mg/kg), oxygen decrease was minimal in both its magnitude and duration, but stronger and more prolonged at higher drug doses (1.0 and 3.0 mg/kg). The rapidity of oxygen response was evident when data were analyzed with rapid, 10-s time resolution for 10 min post-injection (B). In this case, the largest drop in oxygen levels at each dose occurred within 10-30 s from the injection onset, i.e., within the duration of drug delivery.

**Figure 2.**
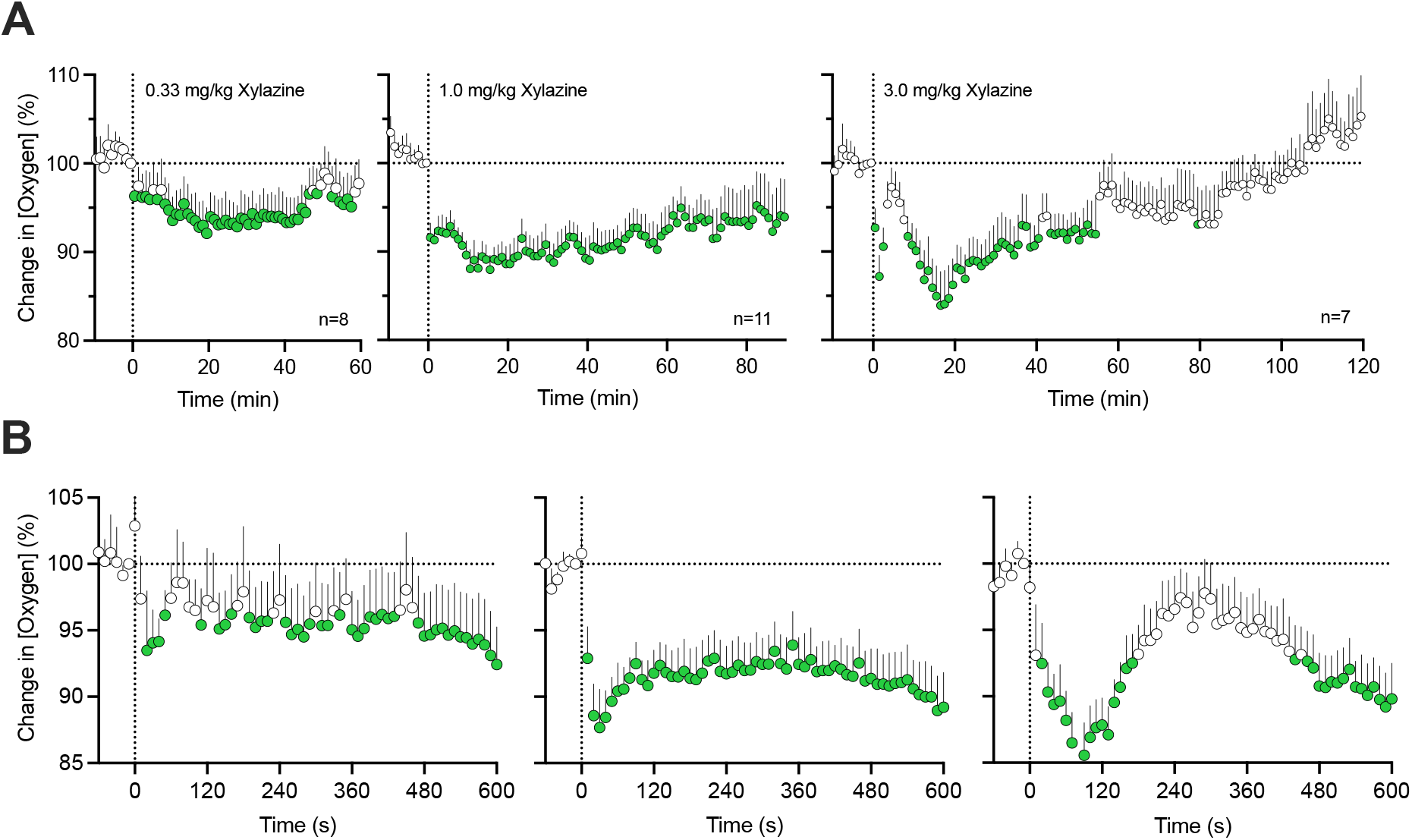
Relative changes in NAc oxygen levels induced by xylazine at different doses (0.3, 1.0 and 3.0) in freely moving rats. A = mean (±SEM) changes assessed with slow (1-min) time resolution. B = mean (±SEM) changes assessed with rapid (10-s) time resolution. Filled symbols show values significantly different from pre-injection baseline. n = numbers of averaged responses

### 3. Effects of xylazine on changes in NAc oxygenation induced by fentanyl

Next, we assessed the effects of a mixture of xylazine at a modest dose (1 mg/kg) on oxygen responses induced by fentanyl (20 μg/kg). These tests were conducted in 5 rats during 16 daily sessions. Consistent with our previous studies [9], fentanyl induced a biphasic NAc oxygen response (F_15,1850_=12.4, p<0.001) with a rapid and strong decrease (~62% below pre-injection baseline) followed by a more prolonged and weaker increase (Figure 3A).

**Figure 3.**
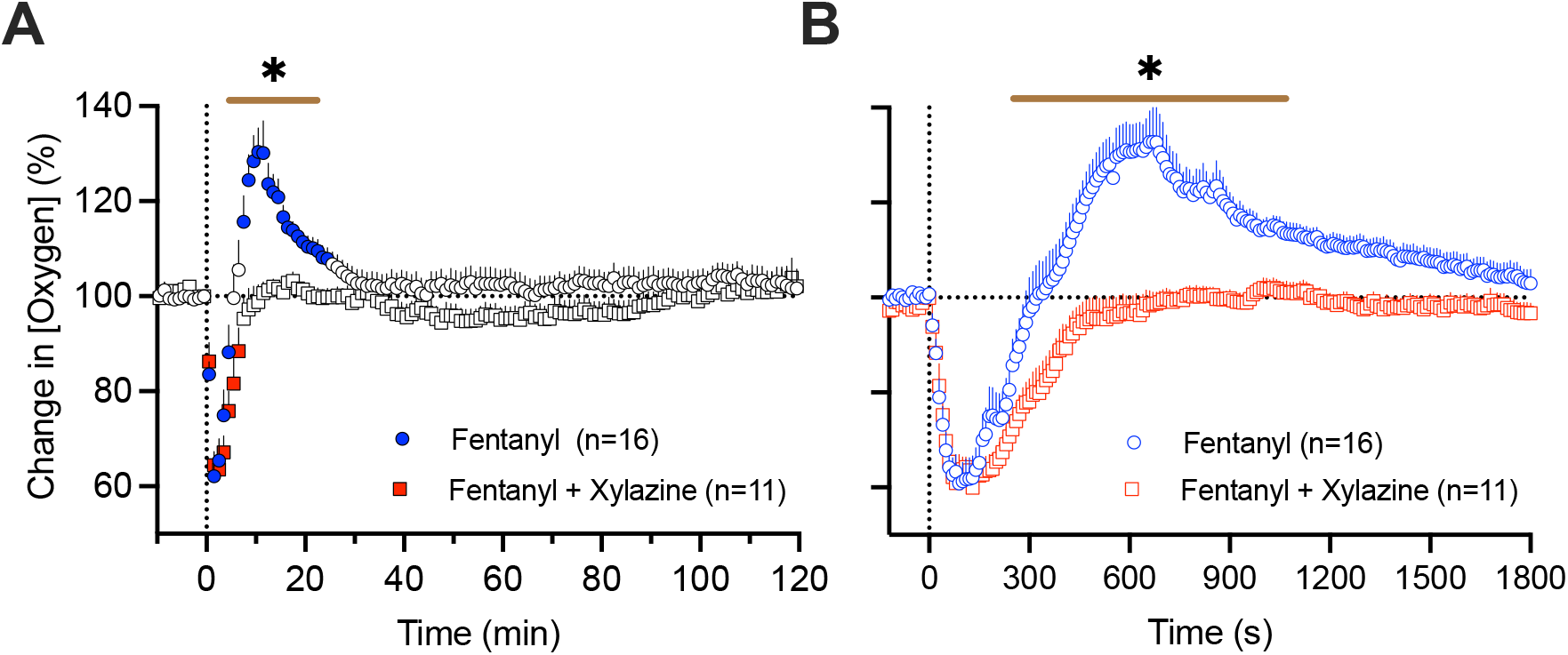
Mean (±SEM) changes in NAc oxygen levels induced by fentanyl (20 ug/kg) and its mixture with xylazine (20 ug/kg + 1 mg/kg) in freely moving rats. A = mean (±SEM) changes assessed with slow (1-min) time resolution. B = mean (±SEM) changes assessed with rapid (10-s) time resolution. Filled symbols show values significantly different from pre-injection baseline. n = numbers of averaged responses. Bold horizontal lines with asterisk show time intervals, during which between-group values were significant.

The xylazine-fentanyl mixture also induced an oxygen decrease (F_10,1210_=12.4, p<0.001), which mirrored the hypoxic response from administering fentanyl alone (~63% below the pre-injection baseline), but lacked the second phase of the oxygen response (Figure 3A). As shown by using a two-way ANOVA with repeated measures, between group-differences were significant from 4 to 22 min (Time x Treatment Interaction F_130,3120_=3.49, p<0.001). Between-group differences were especially evident when the data were analyzed with rapid, 10-s time resolution (Figure 3B). Due to disappearance of the second phase of oxygen response, the total duration of oxygen decrease was longer with the drug mixture than with the fentanyl alone (770 s vs. 330 s). Both fentanyl alone and its mixture with xylazine induced similar behavioral effects, including severe hypoactivity, muscle rigidity in the limbs, tail erection as well as decreases in rate and depth of respiration.

In most cases (10/13), the fentanyl-xylazine mixture induced similar, sedative behavioral effects and the same monophasic pattern of oxygen response (see original examples of oxygen changes induced by fentanyl alone and its mixture with xylazine in Figure 4A and B). However, in three cases (in 2 rats), the xylazine-fentanyl mixture produced convulsions within a couple of minutes post-injection. In this case, oxygen levels strongly decreased but robustly fluctuated in association with convulsion episodes (Figure 4C and D). These three cases were analyzed separately from the main data set.

**Figure 4.**
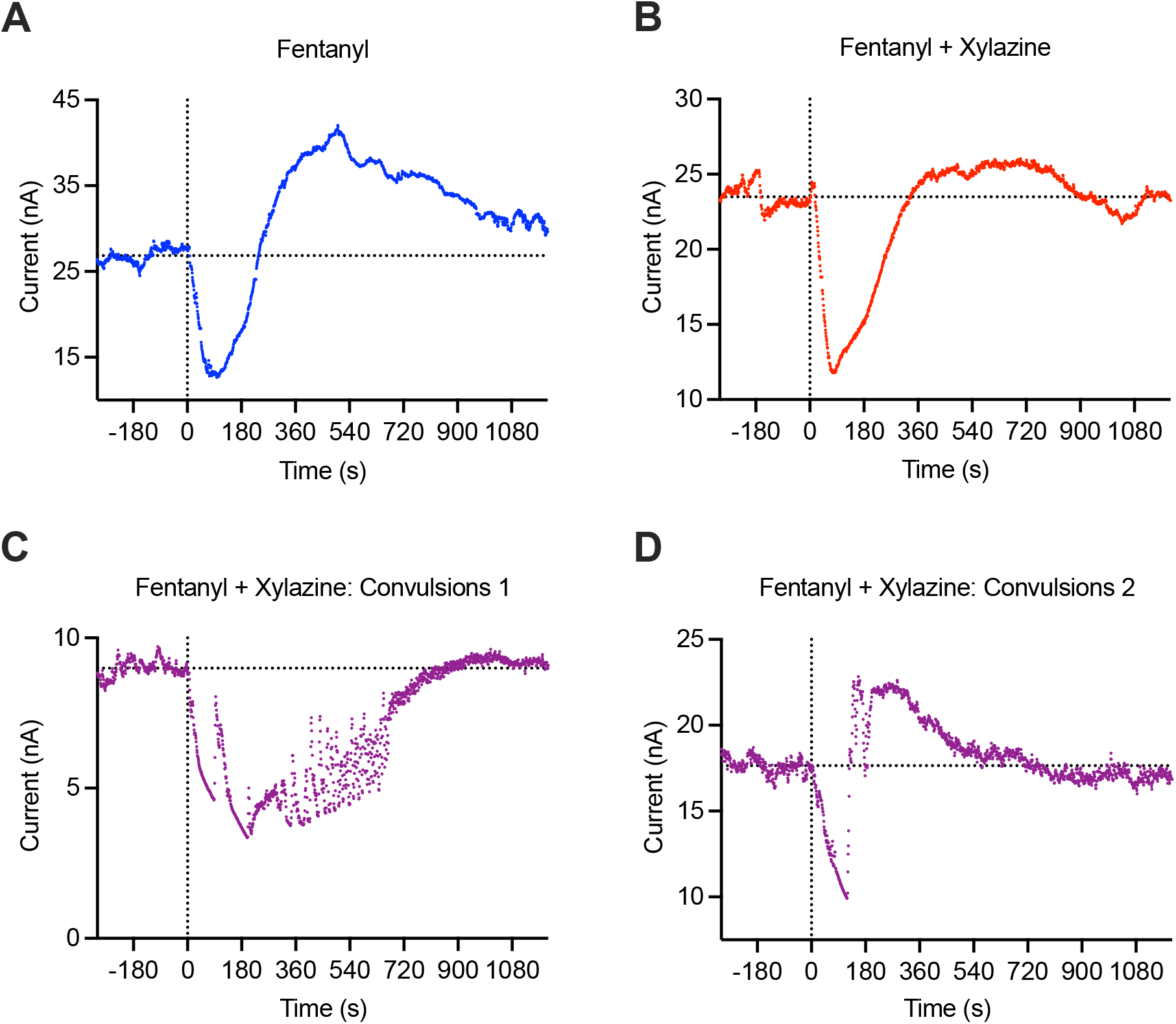
Primary data examples of changes in electrochemical currents (nA) induced by fentanyl and fentanyl-xylazine mixture in freely moving rats. A = fentanyl alone (20 ug/kg), B = fentanyl (20 ug/kg)+xylazine (1 mg/kg), typical example; C and D = unusual changes induced by fentanyl-xylazine mixture with convulsions. Values of reduction current are shown with original (1-s) time resolution, and they were inverted. Since basal reduction currents widely varied between sensors, data were analyzed as the change relative to basal value=100%. Convulsions were never seen after fentanyl alone, but they occurred in 3 cases (in 2 rats) after injections of fentanyl-xylazine mixture.

### 4. Effects of xylazine on changes in NAc oxygenation induced by heroin

The effects of xylazine on NAc oxygenation were tested in 6 rats during 14 daily sessions. As shown in Figure 5, heroin (600 ug/kg) administered alone induced a robust and prolonged decrease in NAc oxygen levels (75.0±3.9 % of baseline for ~17 min) followed by a weaker, more prolonged oxygen increase. The xylazine-heroin mixture also strongly decreased brain oxygenation and this decrease was clearly stronger (61.6±3.0 %; t=2.06, p<0.05) and more prolonged (~72 min) than with heroin alone. As shown by used a two-way ANOVA with repeated measures, the between-group difference was significant (Time x Treatment Interaction F_130,3250_=2.55, p<0.001). Between-group differences in oxygen response were especially evident when we analyzed the areas under the curve for oxygen decrease (4461.1±138.1 vs. 1124±153.6, t=16.2; p<0.0001).

**Figure 5.**
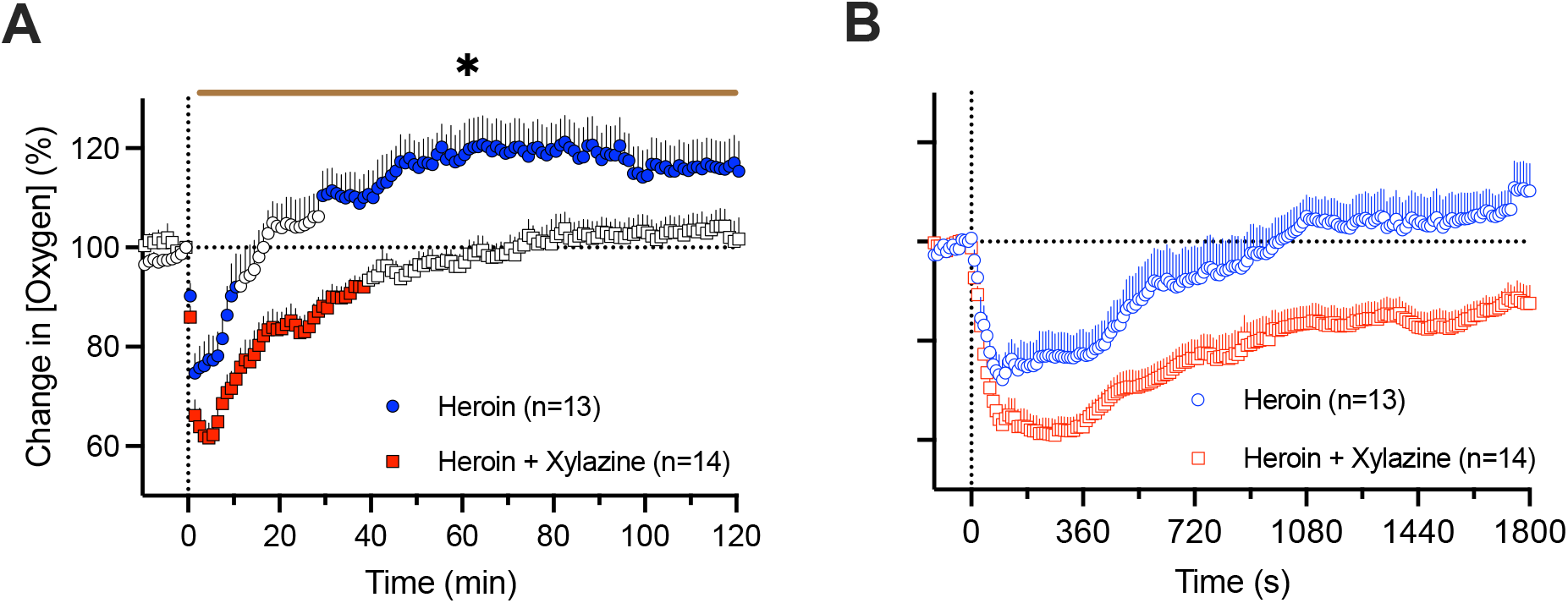
Mean (±SEM) changes in NAc oxygen levels induced by heroin (600 ug/kg) and its mixture with xylazine (600 ug/kg + 1 mg/kg) in freely moving rats. A = mean (±SEM) changes assessed with slow (1-min) time resolution. B = mean (±SEM) changes assessed with rapid (10-s) time resolution. Filled symbols show values significantly different from pre-injection baseline. n = numbers of averaged responses. Bold horizontal lines with asterisk show time intervals, during which between-group values were significant.

## Discussion

We examined the effects of xylazine as an adulterant to highly potent opioids fentanyl and heroin. Since brain hypoxia following respiratory depression is the most dangerous effect of opioid drugs [7, 10, 32, 33], we employed oxygen sensors coupled with high-speed amperometry to examine how xylazine at relatively low, human-relevant doses affects brain oxygenation and whether the addition of xylazine affects the hypoxic effects of fentanyl and heroin, the two drugs largely implicated in overdose-related health complications and death.

Alpha-2 adrenoceptors, the primary substrate for xylazine action, are expressed in both the central and peripheral nervous systems [34–36]. By preferential presynaptic location on central neurons, stimulation of these receptors inhibits the release of norepinephrine, decreasing sympathetic activity and inducing CNS depression. These receptors are also located on smooth muscle cells in blood vessels, and their stimulation induces muscular atonia and decreases vascular tone. Stimulation of these multiple receptors appears to be responsible for a plethora of physiological effects of xylazine, which also depend on the dose and route of administration. In contrast to the relatively large doses of xylazine used during general anesthesia in animals (~10 mg/kg with ip administration in rats), doses used by humans are highly variable and the drug is delivered orally, subcutaneously, and intravenously. The range of toxic doses in humans also greatly varies from 40 to 2400 mg (or 0.6-34.3 mg/70 kg). Since xylazine is rarely used alone and its dosage as adulterant to other more potent drugs is typically low, we chose to test the effects of this drug at relatively low iv doses (0.33-3.0 mg/kg), much lower than the LD50 for iv injections in rats (22-43 mg/kg) [28].

Despite this low-dose exposure, xylazine induced evident sedation, muscle relaxation, and hypothermia—known effects of the drug [37, 38]. In contrast to older studies employing rectal temperature measurements, we used chronically implanted thermocouple sensors, stress-free drug delivery, and high-resolution data analyses to reveal differences in xylazine-induced temperature changes in the brain, temporal muscle, and skin. Although basal temperature in the brain was larger than in temporal muscle, the xylazine-induced decrease was stronger in the muscle than in the brain. This pattern is unusual since temperature changes induced by natural arousing stimuli are typically more rapid and stronger in the brain than in the temporal muscle, suggesting an increase in intra-brain heat production, a sequence of metabolic neural activation [11]. While we could not fully exclude that xylazine may increase metabolic brain activity, stronger temperature decreases in temporal muscle likely result from muscular atonia and atonia-related decreases in heat production [39]. The xylazine-induced temperature decrease in the subcutaneous space was weaker than in the muscle, resulting in a strong increase in the skin-muscle differential. While this change indicates skin vasodilation as a primary reason for heat loss and resulting brain and body hypothermia, we cannot exclude the possibility of an atonia-related decrease in muscular heat production.

Xylazine at large doses can induce respiratory depression and life-threatening hypoxia in animals [40, 41]. Our data revealed that low doses of xylazine delivered iv decreases oxygen levels in the brain, a central effect that may result from respiratory depression. While the effect was weak (93%, 88%, and 82% of baseline levels for 0.33, 1.0, and 3 mg/kg doses), it was prolonged and dose-dependent. This tonic decrease in brain oxygen levels differed from the much stronger but transient decreases induced by fentanyl and heroin, suggesting different underlying mechanisms. These moderate decreases in brain oxygenation may also result from drug-induced decreases in sympathetic activity due to alpha-2 agonism and subsequent tonic decreases in respiratory activity. Alternatively, the weak hypoxic effect of xylazine may be independent of respiratory depression, resulting from cerebral vasoconstriction due to the stimulation of alpha-2 adrenoceptors on cerebral vessels [42].

The known physiological effects of xylazine and its prevalence in opioid-related overdose deaths led us to study the effects of this drug in combination with opioids. The potentiating effect of xylazine and other alpha-2 agonists has been found in early studies on the analgesic effects of opioid drugs, including fentanyl [43]. However, these studies use larger doses of xylazine, and it is still under debate whether the weakening of nociceptive responses reflects the enhancement of opioid analgesia or reinforced sedation by xylazine. To test for the same potentiation effect, we examined xylazine effects at a relatively low, human-relevant dose (1.0 mg/kg) on brain oxygen responses induced by fentanyl and heroin.

We found that the addition of xylazine dramatically changes the pattern of brain oxygen response induced by fentanyl. In contrast to the biphasic effect of fentanyl, with a rapid and strong oxygen decrease followed by its rebound-like increase, the fentanyl-xylazine mixture resulted in elimination of the secondary increase, prolonging the duration of the initial decrease (Fig. 3). Stronger potentiating effects of xylazine were found on decreases in brain oxygenation induced by heroin at a relatively large dose (600 ug/kg; 30:1 ratio vs. fentanyl). Like fentanyl, heroin also induced biphasic oxygen responses, but both the initial decreases and subsequent increases were much more prolonged than with fentanyl. In contrast to heroin alone, the mixture with xylazine induced stronger and more prolonged decreases in brain oxygenation and eliminated the second hyperoxic phase of brain oxygen response.

To determine the possible mechanisms underlying xylazine’s ability to potentiate hypoxic effects of opioid drugs, we need to understand the mechanisms underlying the biphasic changes in oxygen induced by these drugs. In contrast to peripheral tissues, where heroin and fentanyl induce monophasic and relatively prolonged oxygen decreases, the brain hosts two-part oxygen responses with a strong, transient oxygen decrease followed by a weaker, prolonged oxygen increase [26, 44]. While the initial decrease results from respiratory depression and subsequent drop in oxygen in the blood, the subsequent increase may result from cerebral vasodilation and increased vertebral blood flow due to post-hypoxic accumulation of CO2, a powerful vasoconstrictor, in the brain [45–47]. Another factor that mediates cerebral vasodilation induced by opioid drugs is peripheral vasoconstriction that results in redistribution of arterial blood from the periphery to the brain and heart [48]. These two factors likely explain the cerebral vasodilation and increased cerebral blood flow that enhances oxygen entry into brain tissue despite its drop in arterial blood. This is an adaptive mechanism to counteract severe brain hypoxia following insufficient oxygen delivery to the brain from arterial blood.

While the exact mechanisms responsible for prolongation of fentanyl- and heroin-induced hypoxia and disappearance of post-hypoxic hyperoxia induced by xylazine still remain hypothetical, numerous studies point to the vascular effects of this drug in the brain, specifically the blockade of adaptive cerebral vasodilation due to known cerebral vasoconstrictive effects of xylazine and other alpha2 adrenergic agonists [49–51]. Xylazine-induced peripheral vasodilation, which diminishes blood inflow to the brain, and decrease in arterial blood pressure due to xylazine-induced decreased sympathetic outflow may also contribute to the brain-specific hypoxic effects of this drug.

One finding of this study was unexpected. While administration of the fentanyl-xylazine mixture induced sedation in rats during most sessions, the mixture rarely induced convulsions associated with robust fluctuations in brain oxygen levels (Fig. 4). This finding suggests that the effects of the combined use of fentanyl and xylazine are not limited to potentiation of hypoxia and may induce other life-threatening complications. Although the mechanisms underlying this effect of drug combination remain unclear and require further investigation, it is known that convulsions result in robust cerebral vasodilation/increased cerebral blood flow [52, 53], which may counteract the cerebral vasoconstriction induced by xylazine. This explanation remains speculative and require further investigation.

### Conclusions and Human Implications

Overall, our findings deepen the understanding of the involvement of xylazine adulterants in opioid overdose-induced health complications. We highlight the damaging physiological effects of xylazine as it pertains to substance abuse when taken in tandem with opioids by showing the effects of drug mixtures on brain oxygenation and brain temperature. We found that xylazine addition to fentanyl results in elimination of the brain’s compensatory mechanisms to counteract rapid brain hypoxia, and that xylazine addition to heroin potentiates the rapid opioid-induced brain hypoxia in addition to elimination of the following compensatory mechanisms. Our results imply that xylazine can exacerbate the life-threatening potential of opioid use, positing worsened brain hypoxia as a potential cause of death. It will be of clinical importance to observe whether these patterns of oxygenation following xylazine-opioid administration hold and apply in humans.

## Acknowledgements

The study was supported by the Intramural Research Program of the NIH, NIDA

## Author contributions

EAK: Conceptualization, Surgery procedures, Participation in experiments, Data analyses, Writing the manuscript; SC and MRI: Performance of experiments, Data analyses, Graphic work, Histological work, Review and editing the manuscript.

## Data availability

Raw data and the results of their primary analyses are available on request from Dr. Eugene A. Kiyatkin (NIDA-IRP, NIH).

