## Supplemental file for "Xylazine effects on opioid-induced brain hypoxia"

**Table 1**

**Quantitative results of statistical evaluation of temperature responses induced by intravenous injections of xylazine at different doses in freely moving rats**

-------------------------------------------------------------------------------------------------------------

0.33 mg/kg	1 mg/kg	3 mg/kg

Temperature

Change:

NAc	F_(11,671)_=34.35*	F_(11,1001)_=89.99*	F_(10,1210)_=29.66*

Muscle	F_(11,671)_=38.68*	F_(11,1001)_=79.07*	F_(10,1210)_=37.35*

Skin	F_(11,671)_=16.61*	F_(11,1001)_=32.52*	F_(10,1210)_=26.49*

NAc-Muscle

differential	F_(11,671)_=4.313*	F_(11,1001)_=4.968*	F_(11,1210)_=4.308*

Skin-Muscle

Differential	F_(11,671)_=7.122*	F_(11,1001)_=11.95*	F_(11,1210)_=10.25*

n	12	12	11

------------------------------------------------------------------------------------------------------------

The effects of drugs were evaluated with one-way repeated measure ANOVA and F values provide quantitative measure of the effect, with * showing statistical significance with p<001.

n=number of averaged responses.

**Table 2**

**Quantitative results of statistical evaluation of NAc oxygen responses induced by intravenous injections of xylazine at different doses in freely moving rats**

----------------------------------------------------------------------------------------------------------------------

0.33 mg/kg	1 mg/kg	3 mg/kg

1-min time resolution	F_8,488(61)_=10.39*	F_7,637(91)_=7.14*	F_8,968(121)_=10.46*

10-s time resolution	F

n	8	11	7

----------------------------------------------------------------------------------------------------------------------

The effects of drugs were evaluated with one-way repeated measure ANOVA and F values provide quantitative measure of the effect, with * showing statistical significance with p<001.
